# The mutation landscape of *Daphnia obtusa* reveals evolutionary forces shaping genome stability

**DOI:** 10.1101/2025.09.17.676786

**Authors:** Ruyun Deng, Feng Guo, Wen Wei, Michael E. Pfrender, Jeff L. Dudycha, Michael Lynch, Zhiqiang Ye

## Abstract

Spontaneous mutations are the primary source of genetic variation and play a central role in shaping evolutionary processes. To investigate mutational dynamics in *Daphnia obtusa*, we generated a chromosome-level genome assembly spanning 129.4 Mb across 12 chromosomes, encompassing 15,321 predicted protein-coding genes. Leveraging whole-genome sequencing of eight mutation accumulation (MA) lines propagated for an average of 482 generations (spanning over 20 years), we estimated a spontaneous single nucleotide mutation (SNM) rate of 2.23 × 10^-9^ and an indel mutation rate of 2.75 × 10^-10^ per site per generation. The SNM spectrum was strongly biased toward C:G > T:A transitions. Comparative analyses with natural population data revealed that exonic mutations observed in the MA lines were significantly less likely to be present in standing variation than intronic or intergenic mutations, suggesting that purifying selection in natural populations acts to remove deleterious alleles. We also identified 48 *de novo* loss-of-heterozygosity (LOH) events, comprising 8 heterozygous deletions and 40 gene conversion events. The genome-wide gene conversion rate was estimated at 2.62 × 10^-5^ per heterozygous site per generation. These findings provide a comprehensive view of the mutation spectrum, selective pressures, and mechanisms underlying genome stability in *D. obtusa*.

## Introduction

Because spontaneous mutations are the ultimate source of genetic variation, understanding the rate and spectrum of mutations is essential for addressing fundamental questions in molecular evolution, genetic disease, and genome architecture dynamics (Lynch, et al. 2016). Mutation rates influence both the adaptive potential of populations and the accumulation of deleterious load, making them among the most critical parameters in evolutionary biology. However, because *de novo* mutations are rare and often context-dependent, direct measurement requires long-term experimental designs that minimize the confounding effects of natural selection.

Mutation accumulation (MA) experiments, in which organisms are propagated through repeated single-progeny descent to reduce the efficacy of selection, offer a powerful means to estimate spontaneous mutation rates *in vivo* (Halligan and Keightley 2009; Katju and Bergthorsson 2019; Lynch et al. 2023). Nevertheless, most MA studies to date have focused on haploid microbes or obligately sexual species maintained under inbreeding regimes. These systems can exclude mutations such as recessive lethals, large-scale structural variants, and loss-of-heterozygosity (LOH) events. Moreover, the relatively short timescales and limited lineage diversity of many MA studies raise concerns about their ability to capture the full mutational spectrum, particularly for rare and complex mutation types (Baer et al. 2007; Narasimhan et al. 2017; Sasani et al. 2019).

The freshwater microcrustacean genus *Daphnia* is an emerging model for evolutionary and ecological genomics, due to its short generation time, facultative sexual reproduction, and central role in aquatic food webs (Colbourne, et al. 2011; Ebert 2022; Miner et al. 2012). Previous research on *Daphnia pulex* and *Daphnia magna* has revealed relatively high base-substitution mutation rates in both species (Keith et al. 2016; Flynn et al. 2017; Ho et al. 2020). For *D. pulex*, the estimated mutation rates range from 2.3 × 10⁻⁹ to 4.53 × 10⁻⁹ substitutions per site per generation (Keith et al. 2016; Flynn et al. 2017), while the estimate for *D. magna* is somewhat higher, ∼8.9 × 10⁻⁹ per site per generation (Ho et al. 2020). However, the extent to which mutation rates and spectra vary across *Daphnia* species remains poorly understood.

To help fill this gap, we generated a chromosome-level genome assembly of *Daphnia obtusa,* a close relative of *D. pulex* that shares a similar life history and ecological niche—often co-occurring in small, temporary, fishless ponds—it provides an excellent system for comparative analysis due to its well-defined genetic and morphological distinctions. *D. obtusa* is widespread in North American ponds and lakes and reproduces exclusively via cyclical parthenogenesis (Hebert et al. 1987). Morphologically, *D. obtusa* differs from *D. pulex*: females possess large antennular mounds and weak carapace spinulation, while males are characterized by a short postabdominal process and the absence of a postabdominal bay. The presence of hairs on the inner lip of the carapace in females further distinguishes *D. obtusa* from *D. pulex* (Hebert and Finston 1996; Hebert and Crease 1983). Genetically, *D. obtusa* is moderately divergent from the *D. pulex–D. pulicaria* species complex, with a silent-site divergence of ∼0.12, making it highly useful outgroup in population-genomic studies of the latter (Maruki et al. 2022; Ye et al. 2022, 2023). This combination of a shared ecological context with a clear genetic and morphological divergence makes *D. obtusa* an ideal candidate for exploring the evolutionary dynamics of mutation in a comparative framework.

As a foundation for studying mutation and genome evolution in this lineage, we assembled the first chromosome-level reference genome for *D. obtusa*. We then performed deep whole-genome sequencing on eight independently maintained MA lines, each propagated asexually from a single clone for about 500 generations, representing one of the longest-running MA experiments in a multicellular eukaryote. This extensive temporal scale enables the accurate estimation of rare spontaneous mutations, including base substitutions, insertions/deletions (indels), LOH events, and copy number variants (CNVs), as well as the identification of underlying mechanisms such as gene conversion and heterozygous deletion. Finally, by integrating our MA-derived mutational data with genome-wide polymorphism patterns from natural *D. obtusa* populations, we assess the extent to which natural selection shapes mutational outcomes. Together, our results provide a comprehensive view of the mutation and selective landscape in the genome of *D. obtusa*.

## Results

### Chromosome-level genome assembly and functional annotation

To obtain a chromosome-level genome assembly for *D. obtusa*, we used a clone collected from Indiana, USA, and clonally propagated in the laboratory. A total of 12.8 Gb (∼95 ×) PacBio reads were generated and assembled using Canu v1.4 (Koren, et al. 2017). Multiple steps were applied to filter out non-*Daphnia* sequences (potential contamination from food and/or epibionts), e.g., removing contigs with low coverage and abnormal GC content (see Methods). The remaining contigs were scaffolded using 108× reads from high-throughput chromosome conformation capture (Hi-C) technology. The primary draft genome assembly presented here, FS6_V2, consists of 129.4 Mb of DNA sequences located on 12 chromosomes (**Figure 1**). The contig N50 of the assembly is 984 Kb and the scaffold N50 is 11.5 Mb, indicating high contiguity. The BUSCO completeness score (Simão et al., 2015) was estimated at 97.0%, with a duplication rate of 0.6%. The average GC content is 0.41, and approximately 24% of the genome consists of repetitive sequences, such as LTRs, LINEs, as well as DNA transposons and tandem repeats. Among them, LTRs, LINEs, and DNA transposons account for 2.91%, 0.54%, and 0.97% of the genome, respectively.

**Figure 1.**
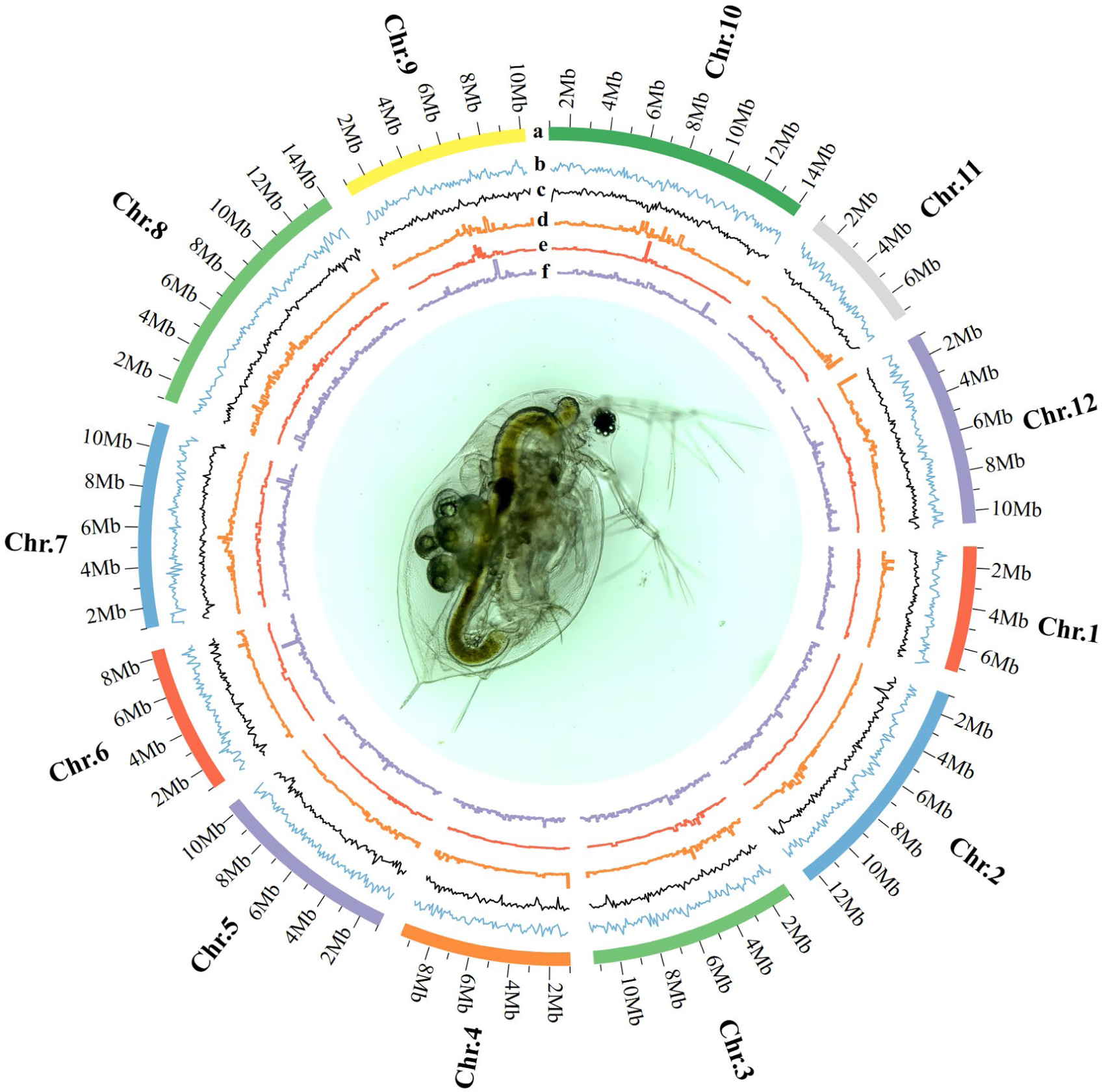
Genomic characteristics of *D. obtusa*. From the outermost to the innermost circle: (a) the 12 chromosomes of the *D. obtusa* genome shown at the megabase (Mb) scale; (b) gene density; (c) GC content; (d) long terminal repeats (LTR); (e) long interspersed nuclear elements (LINE); and (f) DNA transposons, each calculated using a 100 Kb sliding window.

To assist gene prediction, we generated a comprehensive transcriptomic dataset, including 73.1 million Illumina RNA-seq reads and 2.3 million full-length Isoseq reads derived from six environmental stress conditions (high temperature, Atrazine, NaCl, nickel, low pH, and UV light). Using a combined *de novo* and evidence-based annotation strategy (see Methods), we predicted a total of 15,312 protein-coding genes (PCGs) and 5,216 long non-coding RNA genes (**Supplementary Table S1**). Of all PCGs, 94.4% are supported by transcriptomic evidence, whereas only 842 are predicted solely through *ab initio* methods. Among these 842 genes, 384 are also absent in *D. pulex*. Functional annotation was successfully assigned to 11,188 PCGs using Blast2GO, representing 73% of the total.

The genome size of *D. obtusa* is very similar to that of *D. pulex* (129.4 Mb vs. 133.2 Mb). Similarly, the PCGs counts are nearly identical between the two species, with 15,321 in *D. obtusa* and 15,282 in *D. pulex* (**Supplementary Table S1**). In total, 11,230 orthologous genes are shared between them. Notably, the number of predicted long-noncoding RNA genes is slightly higher in *D. obtusa* (5,216 vs. 4,132). In terms of base composition, the GC content is nearly the same (40.9% vs. 40.6%). However, the proportion of repetitive elements is somewhat higher in *D. obtusa* (30.7% vs. 25.7%). Overall, these two species exhibit a high degree of similarity in genome content. The average silent-site divergence between them is 0.12, whereas the average divergence for nonsynonymous sites is 0.013. The synteny map reveals only modest levels of genome rearrangement between the two species (**Supplementary Figure S1**).

### Spontaneous mutation rates and mutational spectra

To estimate mutation rates in *D. obtusa*, we performed deep whole-genome sequencing on eight mutation accumulation (MA) lines. These lines all descend from a single clone collected in November 2001 from a pond in Trelease Woods, Illinois, USA. At the start of the MA experiment, 50 lines were initiated by isolating offspring from a single individual, and these lines were maintained under mutation-accumulation conditions for over 20 years. For this study, eight lines survived for deep whole-genome sequencing, having been propagated asexually for an average of 482 generations (**Table 1**). Each sample yielded an average of 67 million uniquely mapped reads, with a mean coverage breadth of 84.7% and depth of 79× (**Supplementary Table S2**). To ensure high-confidence variant calling, we restricted our analysis to genomic sites with read depths between 20 and 300, resulting in an average of 87 million callable sites per sample (**Supplementary Table S3**).

**Table 1.** Mutation events detected in the MA lines. The number before the hyphen in the sample ID represents the clone, while the number after the hyphen represents the generation number. SNM, single nucleotide mutation; Indel, small (< 20 bp) insertions and deletion.

| Sample ID | Generation | Coverage | SNM | Indel | $\mu(\text{SNM}) \times 10^{-9}$ | $\mu(\text{Indel}) \times 10^{-10}$ |
| --- | --- | --- | --- | --- | --- | --- |
| 13-507 | 507 | 79 | 112 | 6 | 1.26 | 0.68 |
| 14-465 | 465 | 135 | 98 | 3 | 1.17 | 0.36 |
| 20-479 | 479 | 57 | 246 | 33 | 3.08 | 4.13 |
| 26-460 | 460 | 84 | 194 | 27 | 2.40 | 3.33 |
| 27-508 | 508 | 86 | 164 | 21 | 1.83 | 2.34 |
| 28-503 | 503 | 49 | 182 | 18 | 2.23 | 2.21 |
| 33-427 | 427 | 85 | 208 | 24 | 2.76 | 3.19 |
| 48-504 | 504 | 56 | 259 | 48 | 3.11 | 5.77 |
| Average | 482 | 79 | 183 | 23 | 2.23 | 2.75 |

Variants were called using GATK (McKenna *et al*. 2010), and only high-confidence consensus polymorphic sites were retained for downstream analysis. Across the *D. obtusa* genome, 99.43% of sites were homozygous, with only 0.57% being heterozygous. Because all lines were maintained clonally, ancestral heterozygous sites are expected to be preserved unless affected by events such as gene conversion, deletion, or single-nucleotide substitution. We first applied a series of stringent filters to these background heterozygous sites. On average, each line initially contained approximately 0.50 million heterozygous sites, for which we used a binomial test to assess whether allelic ratios conformed to the expected 1:1 distribution. Additional filters included thresholds for minor allele frequency, strand bias, mapping quality, read position quality, and requiring at least five supporting reads for the alternative allele (**see methods**). After filtering, approximately 0.41 million high-confidence background heterozygous sites per line were retained for analysis (**Supplementary Table S3**). After establishing this high-confidence set of background genotypes, we proceeded to identify *de novo* mutations. Based on the assumption that parallel mutations are extremely rare, we defined a *de novo* mutation as a site that was unique to a single MA line and represented a change from the ancestral homozygous state to a heterozygous state.

We identified a total of 1,463 single nucleotide mutations (SNMs) and 180 insertion/deletions (≤5 bp) across all MA lines (**Supplementary File**). Each line harbored between 98 and 259 SNMs and between 3 and 48 indels (**Table 1**). The base substitution mutation rate for each line was calculated as: µ = *x*/(*g* × 2*n*), where *x* represents the number of mutations, *g* represents the number of MA generations, and *n* is the number of callable sites (2*n* accounts for a diploid genome, and the fact that we are identifying heterozygous mutations). The overall SNM rate is estimated to be 2.23 × 10⁻⁹ mutations per base per generation (95% CI: 1.70-2.76 × 10⁻⁹), while the indel rate is 2.75 × 10⁻¹^0^ per base per generation (95% CI: 1.52-3.99 × 10⁻¹^0^). The SNM rate is close to the previous estimates in *D. pulex*, which range from 2.30 × 10⁻⁹ (Flynn et al., 2017) to 4.53 × 10⁻⁹ (for sexual clones, as reported in Table 1 of Keith et al., 2016). Among the base substitutions, the most frequent type is C:G > T:A, occurring at a frequency 5.6 × higher than the least common type, C:G > G:C (**Figure 2A**; **Table 2**). The transition-to-transversion (Ts/Tv) ratio is 1.31 (**Supplementary Table S4**), notably higher than the null expectation of 0.5. This mutational spectrum contrasts with that reported in *D. pulex* by Flynn et al. (2017), who found C:G > A:T to be the most frequent substitution and a Ts/Tv ratio of 0.81. However, our results closely align with those for *D. pulex* reported by Keith et al. (2016), who also observed C:G > T:A as the dominant mutation type and a Ts/Tv ratio of 1.58 (**Figure 2A**), and data from *D. magna* (Ho et al. 2020) reveal a comparable mutational pattern, with C:G > T:A as the most common substitution and a Ts/Tv ratio of 1.54 (**Figure 2A**). We also examined multinucleotide mutations (MNMs), defined as mutations occurring within 50 bp of each other, and identified 93 such events, accounting for approximately 6.4% of all base substitutions. To estimate the expected GC-content of the genome at mutation–equilibrium, we used the rate of G/C → A/T substitutions (*v*) and the rate of A/T → G/C substitutions (*u*). The equilibrium GC-content reflects the long-term balance between GC losses and gains and is calculated as u / (u + v). This model assumes that the nucleotide composition has reached a steady state in which the rate of GC loss equals the rate of GC gain. Based on this calculation, the equilibrium GC content under mutation pressure alone is estimated to be 0.320, which is much lower than the observed GC content of 0.409.

**Figure 2.**
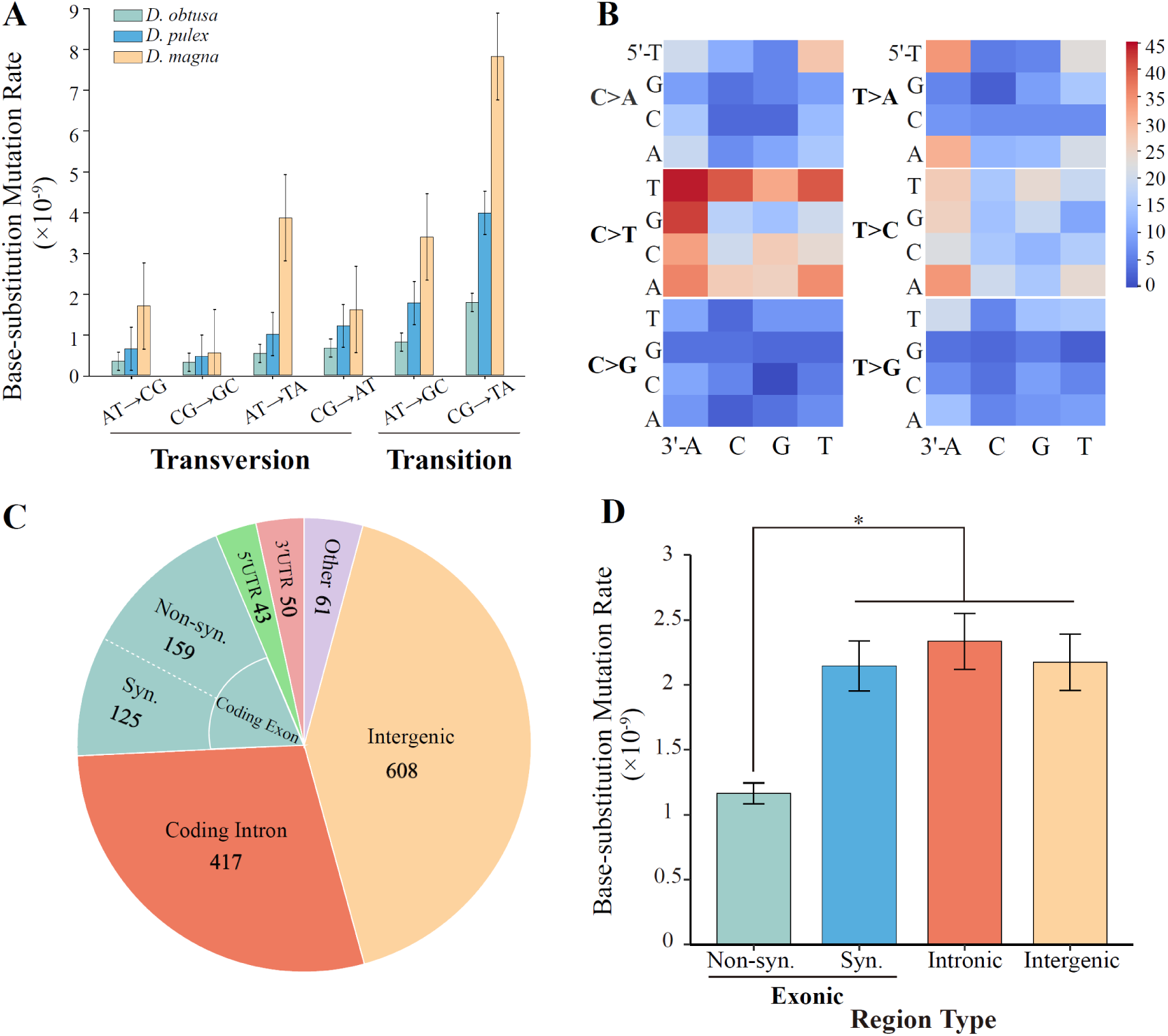
Patterns and distribution of spontaneous mutations in *Daphnia obtusa.* (A) Conditional base mutation rates are compared among *D. obtusa*, *D. pulex* (Keith et al. 2016), and *D. magna* (Ho et al. 2020; averaged across genotypes). (B) The context-dependent mutational spectrum shows the distribution of single-nucleotide mutations by trinucleotide context. (C) Pie chart showing the distribution of mutations across different genomic compartments, including intergenic, exonic, and intronic regions. (D) Mutation rates in intergenic, intronic, and exonic regions are compared. Statistical significance was tested using the χ² test; *p < 0.05.

**Table 2.** Conditional mutation rates of all six possible substitution types.

| Types | Substitution | Frequency | Mutation rate (/site/generation) $\times 10^{-10}$ |
| --- | --- | --- | --- |
| Transition | C:G → T:A | 477 | 17.79 |
| | A:T → G:C | 312 | 8.07 |
| Transversion | A:T → T:A | 206 | 5.31 |
| | C:G → A:T | 178 | 6.62 |
| | A:T → C:G | 132 | 3.40 |
| | C:G → G:C | 85 | 3.14 |

Given the very large mutational harvest, to investigate context-dependent patterns of spontaneous mutations, we analyzed the 96-trinucleotide mutation spectrum, which considers the six types of base substitutions across all 16 possible combinations of flanking 5′ and 3′ nucleotides. We found that substitution rates in *D. obtusa* are significantly influenced by the identity of the central base and its neighboring nucleotides. Specifically, mutations were 1.14 times more frequent when the central base was a C (or G on the complementary strand) than when it was an A or T, highlighting a nucleotide-specific bias. For the most common mutation type, C > T transitions, there is a strong influence of local sequence context. These mutations occurred most frequently when the focal cytosine is preceded by a T at the 5′ position or followed by an A at the 3′ position (**Figure 2B**), suggesting that certain flanking bases enhance mutability at cytosine sites. 5′T–C–A3′ is one of the classic target motifs of APOBEC deaminases, where the flanking 5′T and 3′A context makes the central cytosine more susceptible to deamination into uracil. If this uracil is not repaired before DNA replication, the replication machinery interprets it as thymine (T) and incorporates an adenine (A) opposite it. As a result, the original C:G base pair becomes a T:A base pair in the daughter DNA strand. Over time, this leads to a stable C to T transition mutation in the genome. Interestingly, CpG sites generally show reduced mutation rates, with the exception of C > T transitions, which show elevated mutation frequencies. These results underscore the significant role of local sequence context in shaping the mutational landscape of the genome.

We next assessed mutational bias across different genomic compartments, including intergenic, exonic, and intronic regions. Among the 1,463 SNMs identified, 608 occurred in intergenic regions, 417 in coding introns, and 284 in coding exons (**Figure 2C**). To evaluate whether mutation rates differ among these functional categories, we normalized the mutation counts by their respective genomic proportions. Our analysis revealed that coding introns, intergenic regions, and synonymous sites within exons exhibit a comparable density of accumulated mutations. In contrast, we observed a significant deficit of mutations at nonsynonymous sites (**Figure 2D**; χ2>3, p<0.05 for all comparisons involving nonsynonymous sites). This pattern is not consistent with a uniform accumulation of mutations across all site types. This finding strongly suggests the action of purifying selection, even within the context of our MA experiment. While the experimental design minimizes the efficacy of selection, deleterious nonsynonymous mutations are likely to be removed from the lines, leading to their underrepresentation in the final dataset. Therefore, the reduced frequency of observed nonsynonymous mutations is best interpreted as evidence of strong evolutionary constraint at these sites, rather than a lower intrinsic mutation rate.

To further investigate the potential functional consequences of SNMs, we annotated their predicted impacts using SnpEff (Cingolani et al., 2012). Of all identified SNMs, 2.0% were classified as high-impact variants, typically protein-truncating mutations such as stop-gained or frameshift changes. Approximately 13.9% were predicted to be moderate impact, generally representing missense mutations. Another 8.6% were annotated as low impact, corresponding to synonymous substitutions. The remaining majority (∼75%) were categorized as potential modifiers, referring to variants located in non-coding regions such as introns, untranslated regions (UTRs), or intergenic sequences (**Supplementary Table S5**). To assess whether SNMs are unevenly distributed across chromosomes, we examined their chromosomal distribution. While chromosomes 11 and 12 showed slightly elevated mutation rates, a Chi-square test indicated no significant deviation from a uniform distribution (**Figure S2**; χ² = 3.11, df = 11, p = 0.989).

To investigate how the mutational spectra observed in MA lines compare to those in natural populations of *D. obtusa*, where selection is acting, we sequenced two natural populations—EBG and RAP—each comprising over 90 unique genotypes (**Supplementary Table S6**). Using a minor allele frequency (MAF) threshold of >0.02 to define the presence of alternative alleles, we found that 210 (29%) and 248 (34%) of the SNMs from the MA lines were also present in the EBG and RAP populations, respectively (**Figure 3**), indicating a comparable proportion of missing SNMs across the two populations. Genomic compartment analysis showed that, on average, 32% and 36% of SNMs were present in intronic and intergenic regions, respectively, whereas only 22% and 19% were present in nonsynonymous and synonymous coding sites (**Figure 3; Supplementary Table S7**). The absence of certain SNMs from natural populations can be attributed to several factors, including their absence in the original founding lineages or their subsequent loss through genetic drift or purifying selection.

**Figure 3.**
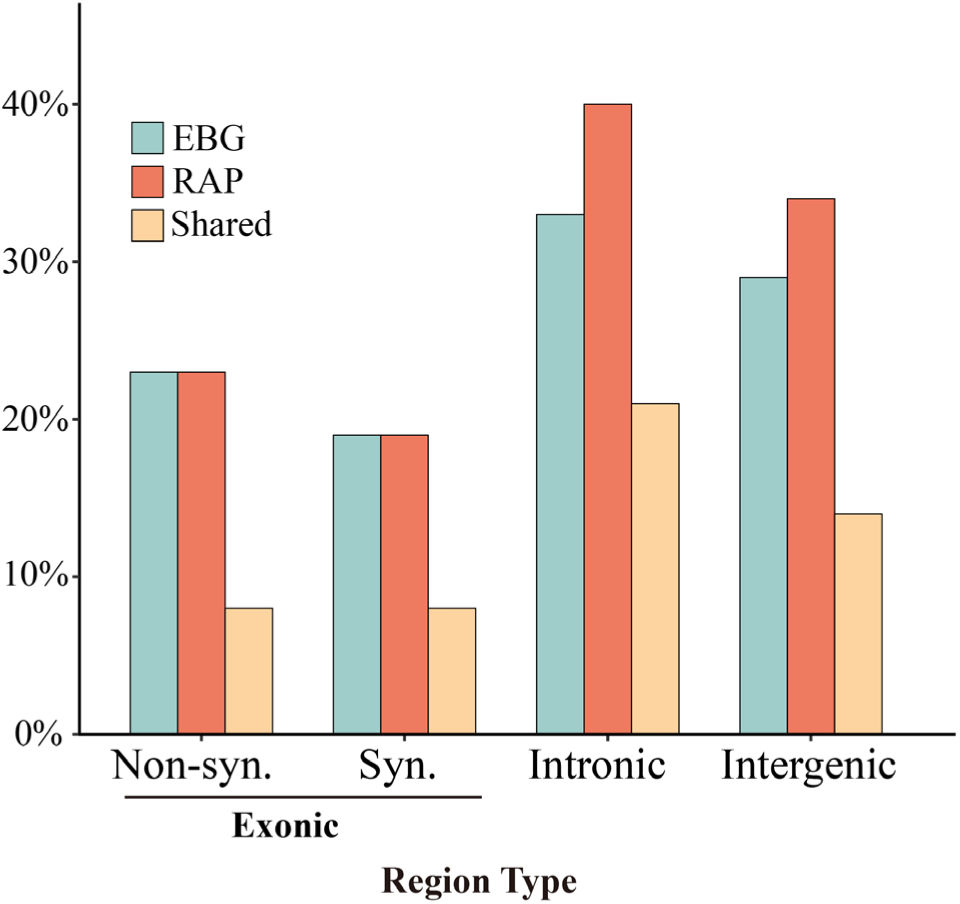
Percentage of single nucleotide mutations (SNMs) observed in the EBG and RAP populations across different genomic categories, including exonic, intronic, and intergenic regions.

Exonic regions are subject to strong functional constraint, meaning mutations within them, particularly nonsynonymous ones, are more likely to be deleterious and removed by direct purifying selection. Additionally, functional regions experience higher rates of both purifying and positive selection, which can reduce local effective population size (Ne) through processes like background selection and selective sweeps. A lower local Ne intensifies the effects of genetic drift, increasing the probability that even neutral or nearly neutral alleles (such as synonymous SNMs) are lost by chance. Therefore, the observed depletion of SNMs in coding regions is consistent with the combined effects of direct selection against deleterious alleles and the indirect effects of linked selection enhancing the power of local genetic drift.

We further estimated the effective population size (*Nₑ*) of the EBG and RAP populations using the formula *Nₑ = πₛ / 4μ*, where *μ* is the mutation rate (2.23 × 10⁻⁹ per site per generation) and *πₛ* is the nucleotide diversity at synonymous sites. Based on *πₛ* values of 0.017 for RAP and 0.012 for EBG, we estimated *Nₑ* values of approximately 1,901,000 and 1,345,000, respectively.

These estimates are substantially higher than those reported for *D. pulex* (*Nₑ* ≈ 640,000; Lynch et al. 2017) and *D. magna* (*Nₑ* ≈ 418,000; Ho et al. 2020), despite *D. obtusa* having a lower mutation rate (2.23 × 10⁻⁹ vs. 2.30–4.53 × 10⁻⁹ in *D. pulex* and 8.9 × 10⁻⁹ in *D. magna*). These relationships are qualitatively consistent with predictions of the drift-barrier hypothesis, which posits that species with larger *Nₑ* tend to evolve lower mutation rates due to more efficient selection against mutator alleles, although such differences can simply arise stochastically as mutation rates drift around the drift barrier (Lynch 2011; Lynch et al. 2016).

### Loss of heterozygosity

Loss of heterozygosity (LOH), defined as the transition from a heterozygous to a homozygous state within a specific genomic region, plays a critical role in evolutionary genetics. LOH can arise via various mechanisms, including gene-conversion-like processes (during mitotic cell divisions, in this case) and heterozygous deletion. To identify *de novo* LOH events, we employed the Runs of Homozygosity (ROH) module implemented in BCFtools (Narasimhan et al., 2016), which applies a hidden Markov model (HMM) to detect homozygous-by-descent tracts based on genotype likelihoods from VCF files. Candidate LOH regions were defined as contiguous tracts longer than 1000 bp with high HMM confidence scores, which consistently contained heterozygous sites in nontarget MA lines. Using this criterion, we identified a total of 48 LOH events, with the number of events per MA line ranging from 0 to 22 (**Table 3; Supplementary File**). The minimal span of these LOH regions ranged from 1.1 kb to 2453.0 kb, while the maximum size range extended from 1.6 kb to 2493.0 kb (**Supplementary File**). Here, the minimum size refers to the distance between the first and last homozygous SNPs within a given LOH event, whereas the maximum size refers to the distance between the first and last homozygous SNPs while also encompassing the flanking heterozygous sites. We estimated a genome-wide LOH rate of 2.93 × 10⁻⁵ per heterozygous site per generation. This estimate is broadly consistent with a previous report for *D. pulex* (4.82 × 10⁻⁵; Flynn et al., 2017), though somewhat lower than that reported by Omilian et al. (2006), who estimated a rate of 1.8 × 10⁻^4^ per heterozygous site per generation. However, direct comparisons across studies should be interpreted with caution due to differences in experimental design and methodological approaches.

**Table 3.** Summary of LOH (loss-of-heterozygosity) events across MA lines. The LOH rate for each mutation accumulation (MA) line was calculated using the formula: µ = *x*/(*g* × *n*), where *x* is the number of LOH sites observed, *g* is the number of MA generations, and *n* is the number of heterozygous sites in the ancestral genotype. Mean LOH tract lengths (the distance between the first and last homozygous SNPs within a given LOH event,) and the number of sites per event are shown with their corresponding standard errors in parentheses. *The total conversion rate was calculated as the average rates across all eight MA lines.

| Sample ID | Events | Mean Length (kb) | Mean Sites | LOH Rates |
| --- | --- | --- | --- | --- |
| 13-507 | 1 | 231 (NA) | 264 (NA) | $1.66 \times 10^{-6}$ |
| 14-465 | 0 | NA | NA | NA |
| 20-479 | 5 | 90 (57) | 248 (143) | $8.36 \times 10^{-6}$ |
| 26-460 | 6 | 404 (205) | 578 (226) | $2.40 \times 10^{-5}$ |
| 27-508 | 1 | 412 (0) | 1595 (0) | $1.02 \times 10^{-5}$ |
| 28-503 | 8 | 397 (302) | 1001 (659) | $5.60 \times 10^{-5}$ |
| 33-427 | 5 | 38 (38) | 164 (140) | $6.05 \times 10^{-6}$ |
| 48-504 | 22 | 338 (85) | 869 (243) | $1.36 \times 10^{-4}$ |
| Total | 48 | 14,324 | 34,538 | $2.93 \times 10^{-5}$* |

To investigate the mechanisms underlying the LOH, specifically whether they result from heterozygous deletions or gene conversion, we analyzed sequencing depth across LOH regions. Regions exhibiting approximately 50% of the sequencing coverage relative to flanking regions and other MA lines were classified as heterozygous deletions. In contrast, LOH regions with coverage similar to their surrounding genomic context were considered consistent with gene conversion-like processes, such as ameiotic recombination. Among the 48 identified LOH events, 8 were attributed to heterozygous deletions, while 40 were attributed to gene conversion (**Supplementary File**). The average length of conversion tracts was 320 kb. The estimated genome-wide rate of conversion-induced LOH was 2.62 × 10⁻⁵ per heterozygous site per generation (**Supplementary Table S8**). This rate is reasonably comparable to that observed in *D. pulex* lines (6.47 × 10⁻⁶; average across obligately asexual and cyclically parthenogenetic *D. pulex* in Keith et al., 2016). However, due to the high variability in the *D. magna* data (Ho et al., 2020), direct comparisons are not feasible.

To explore potential transmission bias, we tracked whether each converted AT/GC heterozygous site was transmitted as an A/T or G/C allele. Converted sites showed a slight excess of GC alleles (50.2%), but this did not deviate significantly from an equal 1:1 ratio (binomial test; p = 0.506), indicating that gene conversion is not biased toward GC. Under this unbiased scenario, the equilibrium GC content is expected to be determined solely by mutation rates, which predict a value of 0.320, yet the observed genome-wide GC content remains stable at 0.409. This discrepancy suggests that forces beyond mutation and conversion—such as selection or other genomic processes—are contributing to the elevated GC content.

## Discussion

This study represents a substantial advance in our understanding of the mutational processes and evolutionary dynamics of *Daphnia obtusa*, and, owing to the extensive mutational data generated, it also provides insights of broader relevance to invertebrates in general. By combining three uniquely valuable genomic resources—a high-quality chromosome-level genome assembly, over two decades of MA data, and population-genomic data from natural populations—we provide an integrative view of how spontaneous mutations arise and are shaped by natural selection. These datasets together form a valuable platform for exploring evolutionary processes in a key aquatic model organism.

Our estimation of the spontaneous mutation rate (μ) in *D. obtusa* falls squarely within the range reported for other eukaryotes, particularly invertebrates (Lynch et al. 2023). We calculated μ to be approximately 2.23 × 10⁻⁹ per site per generation, closely matching prior estimates in *D. pulex* (2.3 × 10⁻⁹ to 4.53 × 10⁻⁹; Keith et al. 2016; Flynn et al. 2017), *Caenorhabditis elegans* (2.7 × 10⁻⁹; Denver et al. 2009), and *Drosophila melanogaster* (2.8– 5.49 × 10⁻⁹; Keightley et al. 2014; Schrider et al. 2013). The observed mutational spectrum, with a strong bias towards C:G > T:A transitions, reflects a conserved pattern found in diverse animals. For instance, a similar transitional bias is a dominant feature of the spontaneous mutation landscape in both humans (Jónsson et al. 2017) and *Drosophila melanogaster* (Schrider et al. 2013). Although this bias is often attributed to methylation and subsequent cytosine deamination, *Daphnia* exhibit minimal DNA methylation largely restricted to gene bodies (Asselman et al. 2015), suggesting that alternative mutagenic processes may be responsible. Moreover, indels occur at approximately one-eighth the frequency of SNPs, are mostly short (<5 bp), and are enriched in repetitive or low-complexity regions—characteristics indicative of replication slippage (Tautz & Schlotterer 1994). Additionally, we observed a Ts:Tv ratio of approximately 1.31, consistent with the universal trend of transitions being more frequent than transversions due to the molecular mechanisms of base misincorporation and repair (Kunkel and Erie, 2005; Lynch et al. 2023).

We next assessed mutational patterns across different genomic compartments and found that, after normalizing by genomic content, mutation rates were broadly similar among intergenic regions, coding introns, and synonymous coding sites. In contrast, nonsynonymous sites exhibited a significant deficit of mutations. This pattern is a classic signature of purifying selection acting even in the context of our MA experiment. Our interpretation is that deleterious nonsynonymous mutations are likely purged from the lines due to their impact on viability, leading to their underrepresentation in the final dataset. Therefore, the reduced frequency of observed nonsynonymous mutations is best interpreted as evidence of strong evolutionary constraint on these protein-coding regions, rather than a lower intrinsic mutation rate. This interpretation is further supported by the functional annotation of the observed mutations. Among all single nucleotide mutations, only ∼2% were classified as high-impact (e.g., stop-gained), 13.9% as moderate-impact (missense), and the vast majority (∼75%) were modifiers in noncoding regions. The scarcity of functionally significant mutations, combined with the pronounced deficit at nonsynonymous sites, paints a cohesive picture: even under MA conditions, strong purifying selection remains an effective filter against highly deleterious mutations. The comparable mutation densities across noncoding and synonymous sites suggest a relatively uniform underlying mutational process, making the depletion at nonsynonymous sites the clearest signal of residual selection during the MA phase.

To investigate the fate of MA-identified mutations in natural populations, we examined their presence in two natural populations (RAP and EBG, each with >90 genotypes). SNMs located in exonic regions—particularly nonsynonymous ones—were less frequently observed than those in intergenic or intronic regions, supporting the hypothesis that purifying selection continues to act in the natural populations, disproportionately eliminating functionally impactful variants. However, interpreting these patterns requires caution. The absence of a given SNM in a natural population does not necessarily indicate selection; it may instead reflect demographic factors, lineage sorting, or the mutation’s absence in ancestral lineages. Interestingly, synonymous SNMs were not retained at a higher rate than nonsynonymous ones, indicating that even silent mutations may be under selection—potentially due to codon usage bias, mRNA stability, or translational efficiency (Chamary & Hurst, 2005; Plotkin & Kudla, 2011).

Finally, we examined the mutation spectrum’s implications for base composition evolution. The mutational spectrum in *D. obtusa* is strongly AT-biased, primarily due to frequent C:G > T:A transitions, which would be expected to reduce genomic GC content over time in the absence of counteracting forces. Although GC-biased gene conversion (gBGC) has been proposed as one such mechanism, our analysis of the observed conversion events revealed no significant GC bias, indicating that gene conversion in *D. obtusa* is effectively unbiased. Under this unbiased scenario, the equilibrium GC content is expected to be determined solely by mutation rates, predicting a value of 0.320, substantially lower than the observed genome-wide GC content of 0.409. This discrepancy suggests that selection, rather than mutation or conversion alone, plays a key role in maintaining elevated GC levels. In particular, selection may favor G/C nucleotides in functionally important regions where they contribute to structural stability, regulatory efficiency, or translational optimization. Thus, while mutation and conversion shape the raw input of genomic variation, natural selection likely acts as a stabilizing force preserving GC content in *D. obtusa*, especially in regions under functional constraint.

## Materials and Methods

### Genome assembly and annotation of *D. obtusa*

We first assembled the PacBio reads using Canu v1.4 (Koren, et al. 2017) with default parameters. Then, multiple filters were applied to the primary assembly: 1) removed contigs with coverage lower than 10 × as they may represent assembly errors; 2) removed contigs with abnormal GC content (<30% or >50%) due to eliminate potential bacterial or algal contamination; 3) performed BLAST searches of each contig against the NCBI Non-Redundant (NR) database and discarded contigs with non-metazoan origins; 4) contigs with low coverage from Hi-C reads (5 ×) were removed from the assembly; 5) flase inclusion of two allelic copies from the heterozygous regions were purged using purge_dups (Guan, et al. 2020); 6) removed mitochondrial contigs. The filtered contigs were then scaffolded into chromosomes using Hi-C interaction data. Hi-C contact maps were generated using Juicer (Durand et al., 2016a) and manually curated with Juicebox v1.11.08 (Durand et al., 2016b).

To estimate genome size, we used a k-mer based approach by creating a histogram from raw Illumina reads with Jellyfish 2.3 (Marçais and Kingsford 2011) and running the GenomeScope web application (http://qb.cshl.edu/genomescope/, last accessed November 2022). RepeatModeler 2.0 (Smit and Hubley 2015) was used to identify *D. obtusa* specific repeats. Repeat sequences in the genome assembly were detected and masked using RepeatMasker-4.1.2-p1 (Smit, et al. 2004) with default setting. Completeness in terms of single copy core orthologs of the final scaffolds was assessed with BUSCO 3.0.2 (Simão et al. 2015), using the Arthropoda set (odb10). (Creation date: 2020-09-10, number of genomes: 90, number of BUSCOs: 1013).

To annotate the genome assembly, we exposed individuals to six abiotic perturbations linked to anthropogenic disturbance. These exposures included: standard and high temperatures (18 °C and 26 °C respectively), low pH (5.0), UV light, nickel (0.03 g/L), Atrazine (4 mg/L), and sodium chloride (5 g/L NaCl). All treatments were maintained for 24 h period without additional algae, except for the UV treatment. The UV exposure was conducted in 250 ml beakers containing 50 ml of medium, placed 10.5 cm below 30 W, 36-inch Reptisun 5.0 UVB fluorescent bulbs for 4 h. All treatments, except high temperature, were conducted at 18 °C, and total RNA was extracted immediately afterward using either the RNeasy Mini Kit or Direct-zol RNA Miniprep Kit (Zymo, Cat. No. R2052). RNA samples were sent to the Translational Genomics Research Institute (TGen) in Phoenix for Illumina sequencing, and to UC Irvine for IsoSeq long-read sequencing. Prior to annotation, repeat sequences were masked as described above. Genome annotation was then carried out using the Eukaryotic Genome Annotation Pipeline - External (https://github.com/ncbi/egapx), which combines *ab initio* and evidence-based gene prediction. Functional annotation of predicted genes was performed using Blast2GO (Götz et al., 2008), allowing assignment of protein domains and Gene Ontology terms. To identify orthologous genes between *D. pulex* and *D. obtusa*, we used orthologous group data from OrthoDB v10 (Kriventseva et al., 2019). Protein sequences for each species were downloaded from NCBI and subsequently clustered into orthologous groups.

### MA line propagating and sequencing

All the MA lines used in this study descend from a single clone of *D. obtusa* collected from a pond in Trelease Woods in Illinois, USA (Latitude: 40.1297; Longitude: −88.1437) in the Spring of 2001. Fifty mutation accumulation (MA) lines were started in November 2001 by taking 50 offspring from a single individual. These 50 offspring became generation 1 of the lines. All MA lines were established and maintained with daily non-quantitative feeding with either *Scenedesmus obliquus* (in Indiana, USA) or *Ankistrodesmus falcatus* (in South Carolina, USA), in beakers with ∼100 mL filtered (1 mm) lake water, and kept in controlled environment chambers at 20°C on a 12:12 L:D photoperiod.

The standardized procedure for propagating the experimental lines was as follows: 8–10 days following the start of the previous generation, a single randomly chosen female offspring was transferred to a new beaker of lake water. Maturation takes place after ∼7 days at 20°, which ensured that the transferred individual was a daughter and not a granddaughter. If a line had not produced offspring by the time of transfer, the mother was transferred to a new beaker and the generation number for that line was not increased. In addition to the focal individual transferred, two of her sisters were transferred into separate beakers to serve as backups. Backups were used for a transfer when the focal individual either died before reproducing or produced only male offspring and/or diapausing eggs over her entire life. Throughout the course of the project, backups were used in ∼10% of the transfers. Use of backups neither showed a trend over time nor was clustered in certain lineages (McTaggart, et al. 2007). Approximately every fifth generation, sisters of the focal individual were collected and frozen at −80 ° for molecular analyses. Several steps were taken to minimize the risk of exogenous contamination and cross-contamination among the lines. Beakers were kept covered to prevent splash contamination when they were not in use. Pipettes used to handle the animals were changed after each transfer to avoid any neonates that may have adhered to the pipette.

Eight lines with generation time close to 500 were survived for sequencing. DNA was extracted from around 100∼500 clonal individuals using the MasterPure™ Complete DNA & RNA Purification Kit (Cat. No. MC85200) following the manufacturer’s protocol. To minimize bacterial contamination, for any DNA collected here for sequencing libraries we starved the culture for 2 d in COMBO media. DNA quality and Concentration was assessed by NanoDrop one and dsDNA HS Qubit Assay (Molecular Probes by Life Technologies, No. Q32851). The integrity of DNA was determined by TapeStation 4200 system. DNA passed quality test was send to The Translational Genomics Research Institute at Phoenix for library construction and sequencing.

### Variants calling and spontaneous mutation rates

To ensure high-quality base-calling, we first removed sequencing adapters and trimmed low-quality bases from the raw reads using Trimmomatic v0.33 (Bolger et al. 2014). The cleaned reads were subsequently mapped to the *Daphnia obtusa* reference genome (FS6_V2) using HISAT2 v2.1.0 (Kim et al. 2019). To retain only uniquely mapped reads, we applied SAMtools v1.8 (Li et al. 2009) with the -q 60 option to exclude those aligning to multiple genomic locations. Variant calling was performed using the HaplotypeCaller module in GATK v4.2.5.0 (Van der Auwera and O’Connor 2020). To ensure high-confidence genotype calls, we applied a stringent filtering pipeline: 1) duplicated reads were removed using Picard MarkDuplicates function (http://broadinstitute.github.io/picard); 2) We excluded SNPs or indels located within 20 base pairs of another indel; 3) tri-allelic sites and multi-allelic sites were excluded from the analysis (∼3% of the total polymorphic sites); 4) Heterozygous sites were retained only if the minor allele frequency (MAF) was ≥ 0.2; 5) only sites with sequencing depth between 20× and 300× were included in downstream analyses; 6) Strand bias was assessed using Fisher’s Exact Test, and sites with bias scores greater than 60 were filtered out; 7) A Mann–Whitney U-based z-approximation was used to test for differences in mapping quality between reads supporting reference and alternate alleles; sites with values less than –12.5 were removed; 8) We also used the Mann–Whitney Rank Sum Test to evaluate read position bias, excluding sites with scores below –8.0; 9) a binomial test was applied to assess deviation from the expected 0.5:0.5 allele balance, and a hard filter was applied to the ALT read counts in the AD field, removing variants with fewer than 5 supporting reads.

To estimate the base substitution mutation rate, we first inferred the ancestral genotype at each polymorphic site, defined as the allele shared by at least seven MA lines. A *de novo* base substitution mutation was identified when one MA line exhibited a novel variant allele while the other lines retained the ancestral genotype. Mutations were excluded if they occurred within 20 bp of an indel or if any of the unmutated lines contained reads supporting the same variant allele, indicating potential pre-existing polymorphisms. Given that over 99% of the *D. obtusa* genome is homozygous, we focused exclusively on Hom > Het mutations, where a homozygous ancestral site becomes heterozygous in the mutated MA line. The nuclear base substitution mutation rate of each MA line was calculated as follows: µ = *x*/(*g* × 2*n*), where *x* represents the number of base substitution mutations that passed the filters, *g* represents the number of MA generations, and *n* is the number of callable sites (2*n* represents diploid genome).

We used the rate of G/C → A/T substitutions (***v***) and the rate of A/T → G/C substitutions (***u***) to estimate the expected GC-content of the genome at mutation–equilibrium. To derive *u* and *v*, we classified all observed base substitutions into two categories: those converting A or T into G or C (AT→GC, contributing to *u*), and those converting G or C into A or T (GC→AT, contributing to *v*). The equilibrium GC-content reflects the long-term balance between GC losses and gains and is calculated using the following formula: GCeq = u / (u + v).

### Mutation spectrum in natural populations of D. *obtusa*

In the spring seasons of 2014 and 2016, we collected *Daphnia obtusa* isolates from seven randomly selected ponds across the United States (**Supplementary Table S6**). To increase the likelihood that each isolate originated from recently hatched resting eggs, adults were sampled early in the growth season. After propagating each isolate clonally in the laboratory, a high mortality rate in populations JP, AQP and RZP resulted in an insufficient number of viable clones for sequencing. For the remaining populations, DNA was extracted from 96 isolates per population. Library preparation was conducted using the Nextera kit, with each sample labeled using a unique oligonucleotide barcode. Sequencing was performed on the Illumina Hiseq 2500 platform, generating paired-end 150 bp reads. To enhance our ability to detect signatures of selection, we aimed to capture genetically diverse individuals by sampling at the time of hatching from sexually produced resting eggs. However, in populations TRH and PYR, most clones appeared to be descendants of the same maternal lineage, as determined by the relatedness function in MAPGD (v0.4.26; Ackerman et al., 2017). This reduced genetic diversity and, consequently, the power to detect selection. Therefore, only populations RAP and EBG, each comprising more than 90 unique genotypes, were retained for downstream analyses.

To prepare data for population-genomic analyses, we constructed nucleotide-read quartets (counts of A, C, G, and T at each site) from raw FASTQ files. After removing adapter sequences with Trimmomatic (v0.36) (Bolger et al. 2014), we aligned the reads to the *D. obtusa* reference genome using Novoalign (v3.02.11) with the “-r None” setting to exclude multi-mapping reads. The resulting SAM files were converted to BAM format using Samtools (v1.3.1) (Li et al. 2009). We then marked duplicates and performed local realignment around indels using GATK (v3.4-0) (McKenna et al. 2010; DePristo et al. 2011; Van der Auwera et al. 2013) and Picard tools. To eliminate redundancy, overlapping read pairs were clipped with BamUtil (v1.0.13), and the final mpileup files were created using Samtools. We generated a comprehensive nucleotide-read profile using the proview command from MAPGD (v0.4.26) (Ackerman et al. 2017).

Despite these preprocessing steps, sequencing artifacts can still affect data accuracy. For instance, some reads may originate from unassembled paralogous regions, leading to anomalous signals at certain sites. To reduce the impact of such mismappings, we excluded any site where a goodness-of-fit test (Ackerman et al. 2017) identified non-binomial read distributions in four or more individuals. We further filtered out clones with mean genome-wide coverage below 3× or clones with total goodness-of-fit scores less than –0.4, which may indicate contamination or other quality issues. To remove potentially unreliable genomic regions, we filtered out sites overlapping with repetitive elements identified using RepeatMasker (v4.0.5) and a custom repeat library (Jurka et al. 2005, last updated August 7, 2022). Additional population-wide coverage thresholds were set to avoid artifacts from under- or over-represented loci. After completing all filtering steps, we estimated allele and genotype frequencies for each population using the maximum-likelihood method implemented in the *allele* function of MAPGD (Ackerman et al. 2017). Finally, sites with error-rate estimates greater than 0.01 were excluded from downstream analysis.

To investigate the mode of natural selection acting on amino-acid sequences of protein-coding genes, we calculated genetic diversity at synonymous (π_S_) and nonsynonymous (π_N_) sites for each gene in both populations, along with the π_N_/π_S_ ratio. For a given site in a specific population, π was estimated as 2*p*(1−*p*), where *p* represents the minor allele frequency obtained from the *allele* function of MAPGD (Ackerman et al. 2017).

### Loss of heterozygosity

To identify loss of heterozygosity (LOH) events in MA lines, we systematically analyzed heterozygous sites across all genotypes. De novo LOH events were detected using the ROH module in BCFtools (Narasimhan et al., 2016), which employs a hidden Markov model (HMM) to identify homozygous-by-descent tracts based on genotype likelihoods from VCF files. Accurate estimation of runs of homozygosity (ROH) in BCFtools requires an external allele frequency (AF) file derived from population-level variant data. We generated this input using the maximum-likelihood method implemented in the allele function of MAPGD (Ackerman et al., 2017). Allele frequencies were estimated from polymorphic sites in the EBG and RAP populations, averaged across the two, and provided as input to BCFtools. VCF files included in the analysis were those from GATK (McKenna et al., 2010).

Candidate LOH regions were defined as contiguous tracts longer than 1000 bp with high-confidence HMM scores. The LOH rate (μ) for each MA line was calculated using the formula: μ = x / (g × n), where *x* is the number of LOH sites observed, *g* is the number of MA generations, and *n* is the number of heterozygous sites in the ancestral genotype. The minimum span of each LOH region was defined as the distance between the first and last sites transitioning from heterozygous to homozygous. The maximum span was defined by including the flanking heterozygous positions.

To distinguish LOH caused by gene conversion from that caused by heterozygous deletions, we examined local sequencing depth. For each MA line, site-specific read depth was standardized by dividing coverage at each site by the mean coverage across all callable sites in the same line. An LOH tract was classified as a putative heterozygous deletion if standardized coverage at the affected region was ≤70% of both the flanking regions and the genome-wide average, while the same region in other lines showed ≥80% standardized coverage relative to the same benchmarks. LOH regions that did not meet these criteria and exhibited coverage comparable to surrounding regions were interpreted as gene conversion events.

### Data accession

The FASTQ files of the raw sequencing data for the *D. obtusa* MA lines are available in the NCBI Sequence Read Archive under accession number PRJNA1272726, and the population sequencing data are available under accession number SAMN18588095 (RAP) and SAMN18588093 (EBG). The *D. obtusa* genome assembly is available in NCBI GenBank under accession number PRJNA1272018 and the corresponding annotation can be found at https://osf.io/a9pf3/files/osfstorage. The *D. pulex* (KAP4) genome assembly is available at NCBI under accession GCF_021134715.1, and its corresponding annotation can be accessed at https://www.ncbi.nlm.nih.gov/genome/annotation_euk/Daphnia_pulex/100/.

## Acknowledgments

We thank Emily Williams (Indiana University), as well as Kevin C. Deitz, Christine Ansell, Christopher S. Brandon, Krista Harmon, and Trenton C. Agrelius (University of South Carolina), for their contributions on maintaining the mutation accumulation lines over the past ∼20 years. This work was supported by the National Natural Science Foundation of China (Grant No. 32471695) and the Fundamental Research Funds for the Central Universities (Grant No. CCNU25JC033) awarded to Z.Y.; the NIH (Grant R35-GM122566-01 to M.L.); the NSF Enabling Discovery through GEnomics (EDGE) program (Grant IOS-1922914 to M.L. and Andrew Zelhof, Indiana University), and NSF IOS-2220696 to M.E.P. and Sen Xu, University of Missouri).

