## Supplementary Tables and Figures for "The mutation landscape of *Daphnia obtusa* reveals evolutionary forces shaping genome stability"

**Table S1.** Comparison of genome assembly and annotation statistics between *D. obtusa* and *D. pulex*.

|  | *D. obtusa* | | | | *D. pulex* | | | |
| --- | --- | --- | --- | --- | --- | --- | --- | --- |
| Chr. | Size (Mb) | # Contig | PCGs | lncRNA | Size (Mb) | # Contig | PCGs | lncRNA |
| Chr.1 | 7.1 | 12 | 871 | 266 | 8.3 | 1 | 1,029 | 200 |
| Chr.2 | 12.9 | 32 | 1,328 | 455 | 13.4 | 2 | 1,325 | 363 |
| Chr.3 | 11.6 | 24 | 1,293 | 453 | 13.3 | 1 | 1,386 | 355 |
| Chr.4 | 9.6 | 19 | 1,002 | 371 | 9.2 | 2 | 969 | 255 |
| Chr.5 | 10.6 | 25 | 1,208 | 429 | 12.0 | 1 | 1,257 | 408 |
| Chr.6 | 8.3 | 15 | 951 | 345 | 7.3 | 1 | 863 | 226 |
| Chr.7 | 11.5 | 18 | 1,711 | 521 | 12.3 | 1 | 1,657 | 446 |
| Chr.8 | 14.4 | 31 | 1,379 | 450 | 14.2 | 2 | 1,428 | 371 |
| Chr.9 | 10.4 | 20 | 1,202 | 477 | 9.6 | 1 | 1,162 | 410 |
| Chr.10 | 14.8 | 23 | 2,109 | 596 | 16.3 | 1 | 2,101 | 495 |
| Chr.11 | 7.0 | 16 | 1,055 | 376 | 6.6 | 1 | 945 | 270 |
| Chr.12 | 11.0 | 22 | 1,203 | 477 | 10.7 | 2 | 1,160 | 333 |
| Sum | 129.3 | 257 | 15,312 | 5,216 | 133.2 | 17 | 15,282 | 4,132 |

**Table S2.** Mapped reads and coverage for the samples. The number before the hyphen in the sample ID represents the clone, while the number after the hyphen represents the generation number. Breadth of coverage is the number of genome bases covered by at least one read divided by the genome size 129,356,989 bp. Depth of coverage refers to the number of times a nucleotide is covered by sequencing reads.

| **Sample ID** | **Reads** | **Breadth of coverage** | **Depth of coverage** |
| --- | --- | --- | --- |
| 13-507 | 66,601,884 | 84.8% | 79 |
| 14-465 | 114,753,386 | 85.5% | 135 |
| 20-479 | 48,315,826 | 84.3% | 57 |
| 26-460 | 70,892,331 | 84.9% | 84 |
| 27-508 | 72,565,846 | 84.9% | 86 |
| 28-503 | 41,518,114 | 84.0% | 49 |
| 33-427 | 72,007,145 | 84.9% | 85 |
| 48-504 | 46,911,072 | 84.0% | 56 |
| Average | 66,695,701 | 84.7% | 79 |

**Table S3.** Summary of genomic site filtering criteria used in this study: (a) sites with read coverage between 20 and 300; (b) heterozygous sites predicted by GATK; (c) sites passing a binomial test for the expected 1:1 allele ratio; and (d) additional filters, including thresholds for minor allele frequency, strand bias, mapping and read position quality, as well as a hard filter removing sites with fewer than five ALT-supporting reads based on the AD field.

| Sample | 20≤x≤300^a^ | GATK^b^ | Binomial test^c^ | Final^d^ |
| --- | --- | --- | --- | --- |
| 13-507 | 88,622,775 | 508,191 | 429,579 | 425,191 |
| 14-465 | 91,015,198 | 519,821 | 425,002 | 422,045 |
| 20-479 | 84,616,425 | 502,945 | 423,115 | 416,038 |
| 26-460 | 89,212,784 | 509,926 | 430,328 | 426,034 |
| 27-508 | 89,388,973 | 497,307 | 418,662 | 414,541 |
| 28-503 | 82,208,355 | 465,554 | 384,926 | 376,442 |
| 33-427 | 89,290,523 | 514,800 | 434,364 | 430,178 |
| 48-504 | 83,730,680 | 459,081 | 383,013 | 376,073 |
| **Average** | **87,260,714** | **497,203** | **416,124** | **410,818** |

**Table S4.** Nuclear substitution spectra (counts) and transition-to-transversion (Ts:Tv) ratios across eight mutation accumulation (MA) lines of *Daphnia obtusa*.

| MALines | CG>TA | AT>GC | CG>GC | CG>AT | AT>CG | AT>TA | Ts:Tv |
| --- | --- | --- | --- | --- | --- | --- | --- |
| 13-507 | 32 | 24 | 8 | 10 | 12 | 20 | 1.12 |
| 14-465 | 31 | 19 | 11 | 15 | 3 | 16 | 1.11 |
| 20-479 | 73 | 54 | 8 | 37 | 27 | 33 | 1.21 |
| 26-460 | 65 | 44 | 12 | 22 | 11 | 31 | 1.43 |
| 27-508 | 52 | 34 | 16 | 23 | 16 | 21 | 1.13 |
| 28-503 | 67 | 42 | 4 | 17 | 16 | 27 | 1.70 |
| 33-427 | 77 | 43 | 16 | 25 | 15 | 24 | 1.50 |
| 48-504 | 80 | 52 | 10 | 29 | 32 | 34 | 1.26 |
| Total/Mean | 477 | 312 | 85 | 178 | 132 | 206 | 1.31 |

**Table S5**. Summary of variant effects by impact level, as estimated by SnpEff (Cingolani et al., 2012).

| Type | Count | Percent |
| --- | --- | --- |
| High | 15 | 2.0% |
| Moderate | 102 | 13.9% |
| Low | 63 | 8.6% |
| Modifier | 552 | 75.4% |

**Table S6.** summarizes the *D. obtusa* population collected in this study. EBG and RAP are the ones used the check the power of selection in natural clones.

| Population | Long name | State (USA) | Latitude, Longitude | Year | Live clones |
| --- | --- | --- | --- | --- | --- |
| **EBG** | **Edinburgh** | **MO** | **40.0814, -93.6938** | **2015** | **130** |
| PYR | Pyramid Court | IN | 39.2099, -86.5793 | 2014 | 108 |
| AQP |  | AL | 33.0584,-87.6406 | 2016 | 72 |
| JP |  | GA | 33.5658,-85.0872 | 2016 | 9 |
| **RAP** | **Refuge Admin** | **MS** | **33.2691,-88.8088** | **2016** | **119** |
| RZP |  | AR | 33.8089,-92.8897 | 2016 | 46 |
| TRH |  | AL | 33.0584,-87.6404 | 2016 | 116 |

**Table S7.** Summary of SNMs by functional category. *Shared* denotes variants found in both EBG and RAP.

| Region Type | Total | EBG | RAP | Shared |
| --- | --- | --- | --- | --- |
| Exonic | 153 | 33 | 33 | 12 |
| synonymous | 48 | 9 | 9 | 4 |
| nonsynonymous | 105 | 24 | 24 | 8 |
| Intronic | 167 | 49 | 57 | 24 |
| Intergenic | 283 | 92 | 113 | 59 |

**Table S8.** Summary of gene conversion level events across MA lines.
The conversion rate for each MA line was calculated using the formula: µ = *x*/(*g* × *n*), where *x* is the number of conversion sites observed, *g* is the number of MA generations, and *n* is the number of heterozygous sites in the ancestral genotype. Mean conversion tract lengths and the number of sites per event are shown with their corresponding standard errors in parentheses. Total indicate the number summarized across all lineages.

| **Sample ID** | **Events** | **Mean Length (kb)** | **Mean Sites** | **Conversion Rates** |
| --- | --- | --- | --- | --- |
| 13-507 | 0 | NA | NA | NA |
| 14-465 | 0 | NA | NA | NA |
| 20-479 | 3 | 14 (7) | 60 (53) | 8.36×10-6 |
| 26-460 | 6 | 404(205) | 578 (226) | 2.40×10-5 |
| 27-508 | 1 | 412 (NA) | 1595 (NA) | 1.02×10-5 |
| 28-503 | 7 | 452(343) | 1136 (745) | 5.60×10-5 |
| 33-427 | 5 | 385(34) | 164 (140) | 6.05×10-6 |
| 48-504 | 18 | 363(102) | 935 (284) | 1.36×10-4 |
| Total | 40 | 14,762 | 30,868 | 2.62×10^-5*^ |

**Table S9**. Duplication and deletion rates for de novo CNVs >1 kb across MA lines. The base-level mutation rate (μ_CNV_) was calculated using the formula: μ_CNV_ = n / (2 × G × N), where *n* is the number of de novo CNV events (>1 kb), *G* is the total number of generations, and *N* is the number of sites with coverage between 20 and 300 (see Table S3).

| Sample ID | deletion | duplication | μ (del.) | μ (dup.) |
| --- | --- | --- | --- | --- |
| 13-507 | 54 | 1 | 6.01×10^-10^ | 1.11×10^-11^ |
| 14-465 | 73 | 0 | 8.62×10^-10^ | NA |
| 20-479 | 85 | 0 | 1.05×10^-9^ | NA |
| 26-460 | 5 | 113 | 6.09×10^-11^ | 1.38×10^-9^ |
| 27-508 | 6 | 17 | 6.61×10^-11^ | 1.87×10^-10^ |
| 28-503 | 41 | 27 | 4.96×10^-10^ | 3.26×10^-10^ |
| 33-427 | 8 | 48 | 1.05×10^-10^ | 6.29×10^-10^ |
| 48-504 | 37 | 16 | 4.38×10^-10^ | 1.90×10^-10^ |
| Average | 39 | 28 | 4.60×10^-10^ | 3.40×10^-10^ |

**Figure S1. Synteny map between *Daphnia obtusa* and *Daphnia pulex*.**
The map was generated by comparing the whole-genome assemblies of *D. obtusa* and *D. pulex* using the online tool [D-GENIES](https://dgenies.toulouse.inra.fr).

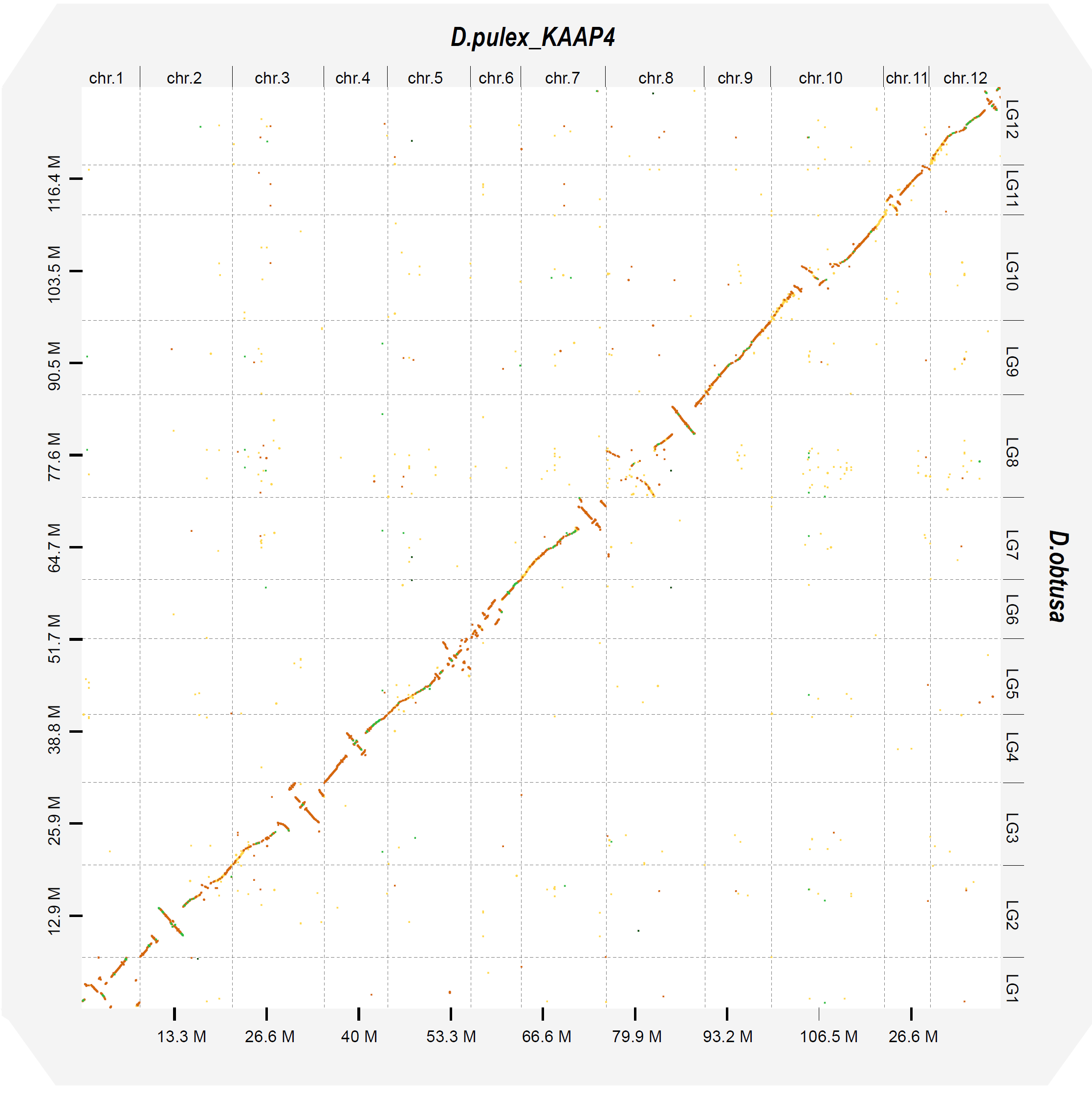

**Figure S2**. Mutation rates across the 12 chromosomes reveal no statistically significant differences.

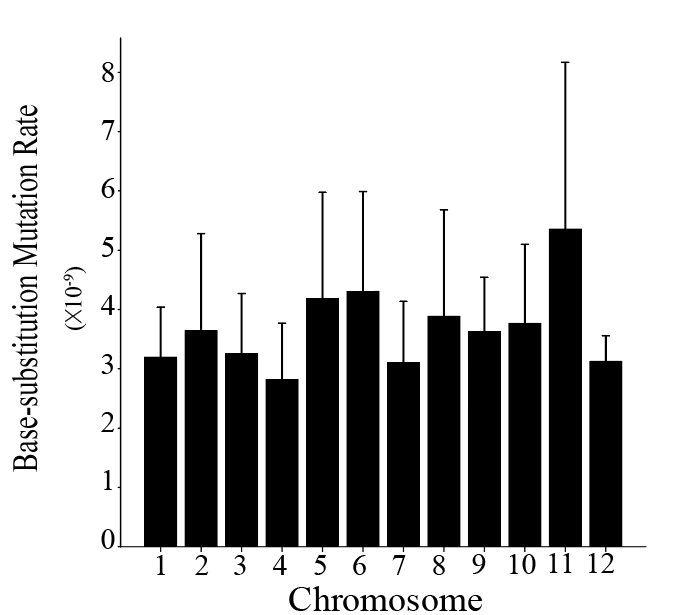
